# Automated identification of Cell Types in Single Cell RNA Sequencing

**DOI:** 10.1101/532093

**Authors:** Feiyang Ma, Matteo Pellegrini

## Abstract

Cell type identification is one of the major goals in single cell RNA sequencing (scRNA-seq). Current methods for assigning cell types typically involve the use of unsupervised clustering, the identification of signature genes in each cluster, followed by a manual lookup of these genes in the literature and databases to assign cell types. However, there are several limitations associated with these approaches, such as unwanted sources of variation that influence clustering and a lack of canonical markers for certain cell types. Here, we present ACTINN (Automated Cell Type Identification using Neural Networks), which employs a neural network with 3 hidden layers, trains on datasets with predefined cell types, and predicts cell types for other datasets based on the trained parameters. We trained the neural network on a mouse cell type atlas (Tabula Muris Atlas) and a human immune cell dataset, and used it to predict cell types for mouse leukocytes, human PBMCs and human T cell sub types. The results showed that our neural network is fast and accurate, and should therefore be a useful tool to complement existing scRNA-seq pipelines.

**Author Summary:** Single cell RNA sequencing (scRNA-seq) provides high resolution profiling of the transcriptomes of individual cells, which inevitably results in high volumes of data that require complex data processing pipelines. Usually, one of the first steps in the analysis of scRNA-seq is to assign individual cells to known cell types. To accomplish this, traditional methods first group the cells into different clusters, then find marker genes, and finally use these to manually assign cell types for each cluster. Thus these methods require prior knowledge of cell type canonical markers, and some level of subjectivity to make the cell type assignments. As a result, the process is often laborious and requires domain specific expertise, which is a barrier for inexperienced users. By contrast, our neural network ACTINN automatically learns the features for each predefined cell type and uses these features to predict cell types for individual cells. This approach is computationally efficient and requires no domain expertise of the tissues being studied. We believe ACTINN allows users to rapidly identify cell types in their datasets, thus rendering the analysis of their scRNA-seq datasets more efficient.

## Introduction

Single cell RNA sequencing (scRNA-seq) enables the profiling of the transcriptomes of individual cells, thus characterizing the heterogeneity of samples in manner that was not possible using traditional bulk RNA-Seq^[1]^. However, scRNA-seq experiments typically yield high volumes of data, especially when the number of cells is large (often many thousands). Thus, fast and efficient computational methods are essential for scRNA-seq analyses.

One common goal of scRNA-seq analyses is to identify the cell type of each individual cell that has been profiled. To accomplish this, typically cells are first grouped into different clusters in an unsupervised way, and the number of clusters allows us to approximately determine how many distinct cell types are present in the sample. Each cluster should contain cells with similar expression profiles, and so the aggregated profile of a cluster increases the signal to noise of the expression estimates. To attempt to interpret the identity of each cluster, marker genes are found as those that are uniquely highly expressed in a cluster, compared to all the other clusters. These canonical markers are then used to assign the cell types for the clusters, by cross referencing the markers with lists of previously characterized cell type specific markers. While this process is able to identify cell types, there are some limitations: 1. Since the clustering method is unsupervised, all sources of variation influence the formation clusters, including effects that are not directly related to cell types such as differential expression induced by cell cycles. 2. It is often difficult to find an optimal match between the marker genes associated with each cluster and the canonical markers for specific cell types. Moreover, depending on the clustering parameters used, one cluster might contain multiple cell types, or one cell type could be split into multiple clusters. 3. Using canonical markers to assign cell types requires background knowledge of cell type specific markers, and sometimes these are not well characterized or difficult to find in the literature. Moreover, some canonical markers may be expressed by more than one cell type, and some cell types may have no known markers. 4. The same types of cells processed by two distinct scRNA-seq techniques tend to cluster separately due to technical batch effects, which complicates cell type identification in composite datasets. 5. Cell subtypes are often very similar to each other, which limits efforts to separate them accurately into different clusters. To overcome many of the limitations of existing approaches, new methods need to be developed.

Neural networks provide a popular framework for machine learning algorithms which can be used to interpret complex datasets. As a result, neural networks have been widely used in many fields, including for the analysis of scRNA-seq data^[2-5]^. Since the output data from scRNA-seq is feature-enriched and well-structured, it is well suited as an input for neural networks. Here, we present ACTINN (Automated Cell Type Identification using Neural Networks) for scRNA-seq cell type identification. To overcome may of the limitations of traditional cell type identification approaches described above, we used a neural network with 3 hidden layers, trained it on scRNA-seq datasets with predefined cell types, and predicted cell types in other datasets based on the trained parameters. We tested our neural network with several published datasets and show that it is fast, efficient and accurate.

## Results

### Overview of the neural network

We used a neural network with 3 hidden layers, each containing 100, 50 and 25 nodes, respectively (Fig 1). For the activation functions, we used the softmax function for the ouput layer and the rectified linear unit (ReLU) function for the other layers. We used the cross-entropy function as the loss function. The neural network model was implemented using TensorFlow (https://www.tensorflow.org/), and the code was written in python. We trained the neural network on 6 Intel(R) Xeon(R) CPU E5-2660 v3, and the training process took 0.5 minute to complete with 1000 cells, 11 minutes with 32,000 cells and 21 minutes with 56,000 cells. The maximum memory used in training with 56,000 cells was 18 GB. The code and datasets used in this study are available at https://github.com/mafeiyang/ACTINN.

**Fig 1.**
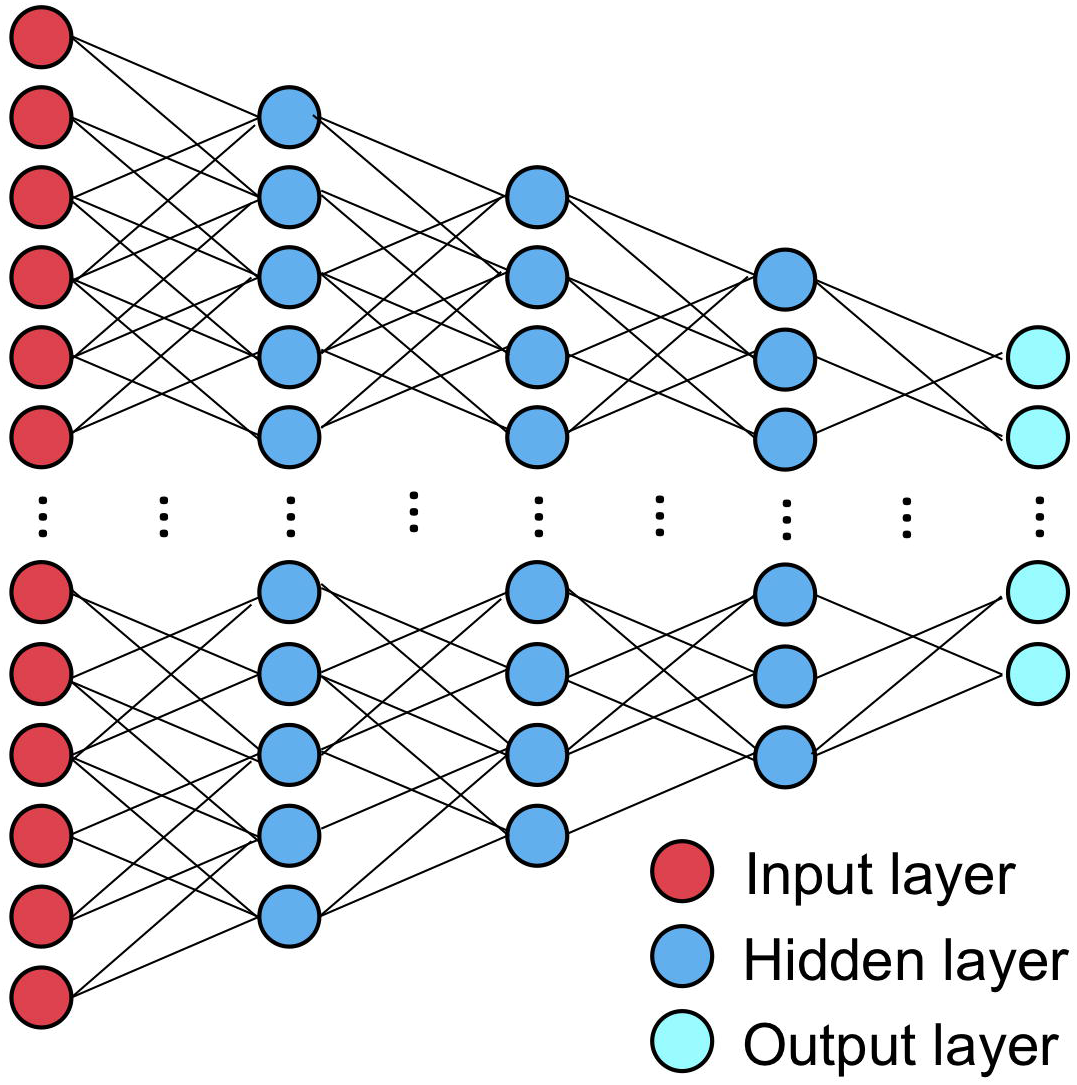
Neural network configuration.

### ACTINN model for murine cell types

We used 2 datasets from the Tabula Muris Consortium (The Tabula Muris Consortium. 2018) to train and test our neural network. The datasets contain 100,605 cells from 20 mouse organs, and were sequenced by two distinct techniques, 10X Genomics (10X) and Smart-seq2 (SS2). To ensure we are using cells with high quality, we filtered out cells with less than 300 detected genes, clustered the cells, and identified marker genes for each cluster using Seurat^[6]^. The details of the Seurat analysis can be found in the methods section. We manually assigned cell types for each cluster based on canonical markers (Fig 2A). We focused on 12 cell types and selected cells that have the same labels between our analyses and the Tabula Muris Consortium’s. This process resulted in 56,112 cells (Fig 2B). Cells processed by 10X have a median of 4,787 unique molecular identifiers (UMIs) and 1,558 genes detected, and cells processed by SS2 have a median of 623,799 UMIs and 2,448 genes detected.

**Fig 2.**
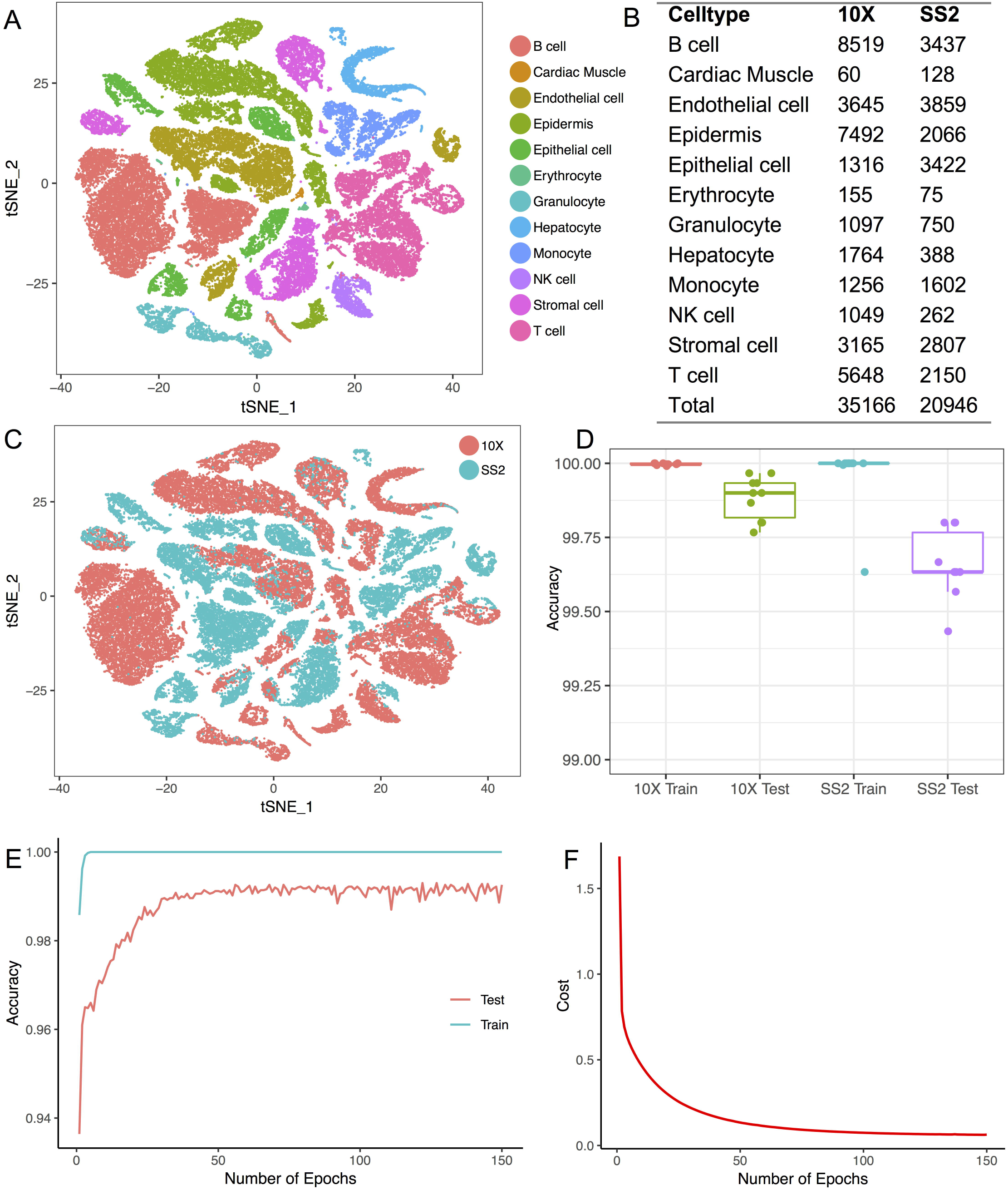
Training and testing of the neural network on the Tabula Muris Atlas. (a) Cell types obtained from the TMA. (b) Number of cells obtained for each cell type from each technique. (c) The same cell type tends to cluster separately by techniques. (d) Training and testing accuracy of the neural network when trained and tested using cells processed by the same technique. (e) Training and testing accuracy after each epoch when trained with 5,000 10X cells and tested with 5,000 SS2 cells. (f) Cost after each epoch when trained with 5,000 10X cells and tested with 5,000 SS2 cells.

To test the robustness of our neural network’s performance, we first trained and tested it on cells processed by each scRNA-seq platform separately. To this end, we randomly sampled 3000 cells for testing, and used the remainder of cells for training. We repeated this process 10 times, and the average training accuracies for the 10X dataset and the SS2 dataset were 99.997% and 99.963%, respectively, and the average testing accuracies were 99.883% and 99.660%, respectively (Fig 2D). These results show that our neural network can achieve very high accuracy when training and testing on datasets generated by the same technique.

### ACTINN overcomes batch effects introduced by different techniques

Different scRNA-seq techniques can introduce significant batch effects^[7]^ with the same cell types clustering separately due to technical artifacts (Fig 2C). To test our neural network’s performance accounting for the batch effects introduced by different techniques, we trained it on cells processed by one platform and tested it on cells processed by the other. We first trained the neural network on all the 10X cells and tested in on all the SS2 cells. The training accuracy was 99.997% and the testing accuracy was 98.625%. Among the 288 incorrectly predicted cells, 118 monocytes were predicted as B cells, 64 monocytes were predicted as epithelial cells, 47 NK cells were predicted T cells (Supplementary File 1). We then trained the neural network on the SS2 dataset and tested it on the 10X dataset. The training accuracy was 100% and the testing accuracy was 99.195%. Among the 283 incorrectly predicted cells, 150 endothelial cells were predicted as epidermis, 46 T cells were predicted as NK cells, and there were several other mispredictions (Supplementary File 1).

### Early stopping prevents overfitting of the training set

To prevent overfitting the parameters on the training set, we randomly sampled 5,000 cells from the 10X dataset and 5,000 cells from the SS2 dataset. We trained the neural network on the 10X cells and tested it on the SS2 cells. During the training process, we recorded the accuracy and the cost after each epoch. The accuracy was defined as the percentage of cells whose cell type was correctly predicted, and the cost was the output of the cost function after each epoch. We found that the training accuracy saturated early (5 epochs), and the testing accuracy saturated at around 50 epochs (Fig 2E), and the cost decreased very slowly after 50 epochs (Fig 2F). These results indicate that early stopping can be used to reduce training time and prevent overfitting.

### Cell type prediction using the mouse cell atlas

Since the cell types from the two mouse cell atlas datasets can be accurately predicted, we combined the two datasets and used the combined dataset as the reference to predict cell types for other datasets. We first tried to predict cell types for a dataset that contains flow cytometry sorted leukocytes from mouse aorta^[8]^. All cells were predicted as leukocytes except for 1 erythrocyte, which we think is a doublet of an erythrocyte and B cell as high expression of hemoglobin genes was detected (Fig 3A). We also carried out unsupervised analysis on the dataset and clustered the cells using Seurat. Then we used the canonical markers to assign the cell types for each cluster (Fig 3B). Most cells had the same cell type assignment by the two methods. However, our neural network detected some natural killer (NK) cells, which were in the same cluster with the T cells, and were assigned as T cells in the unsupervised clustering. We checked the expression of CD3D, CD8A and GZMA (Fig 3C), and found no expression of CD3D and CD8A, but high expression of GZMA in the NK cells, which suggests that these are likely NK cells.

**Fig 3.**
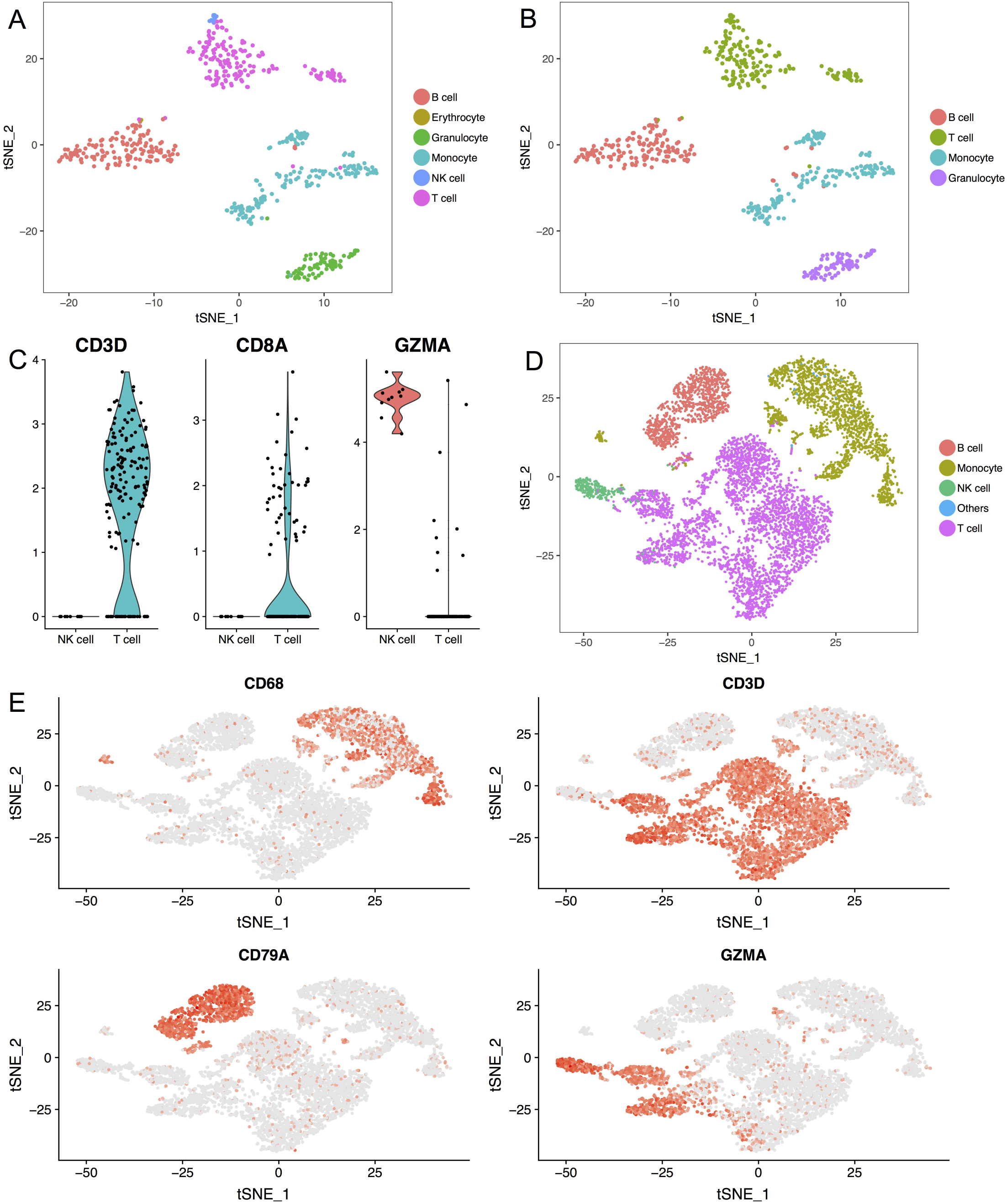
Neural network predicts cell types for human and mouse datasets. (a) Cell types predicted by the neural network for the mouse leukocyte dataset. (b) Cell types identified by unsupervised clustering and canonical markers for the mouse leukocyte dataset. (c) Violin plots showing 3 genes’ expression level in the NK and T cells from the mouse leukocytes. (d) Cell types predicted by the neural network for the human PBMC dataset. (e) TSNE plots showing 4 marker genes’ expression for the human PBMC dataset.

It is generally thought that human and mouse share similar cell types, and the same cell type from human and mouse share similar expression profiles. To test this, we trained our neural network on the mouse cell atlas datasets and used the parameters to predict the cell types for a human peripheral blood mononuclear cell (PBMC) dataset. We found 4 main populations in the PBMC dataset, namely, B cells, monocytes, NK cells and T cells (Fig 3D). We plotted the canonical markers for these 4 populations (Fig 3E) and found that the predicted cell types matched the expected marker expression. These results suggest that the mouse cell atlas datasets can be used as a reference to identify cell types for both human and mouse cells.

### ACTINN accurately predicts cell subtypes

Although it is relatively easy to distinguish different cell types in scRNA-seq using the unsupervised clustering methods, it is more difficult to further divide one cell type into cell subtypes. Here, we collected 5 publicly available datasets^[9]^, each containing one flow cytometry sorted T cell subtype. We merged these datasets and selected the cells that have the same labels between our analyses and the flow cytometry sorting, and then used these cells as a reference for the neural network. We then clustered the selected cells and identified markers (Fig 4A and 4B) for each sub cell type using Seurat. For the test set, we used the T cells from the human PBMC datasets mentioned above.

**Fig 4.**
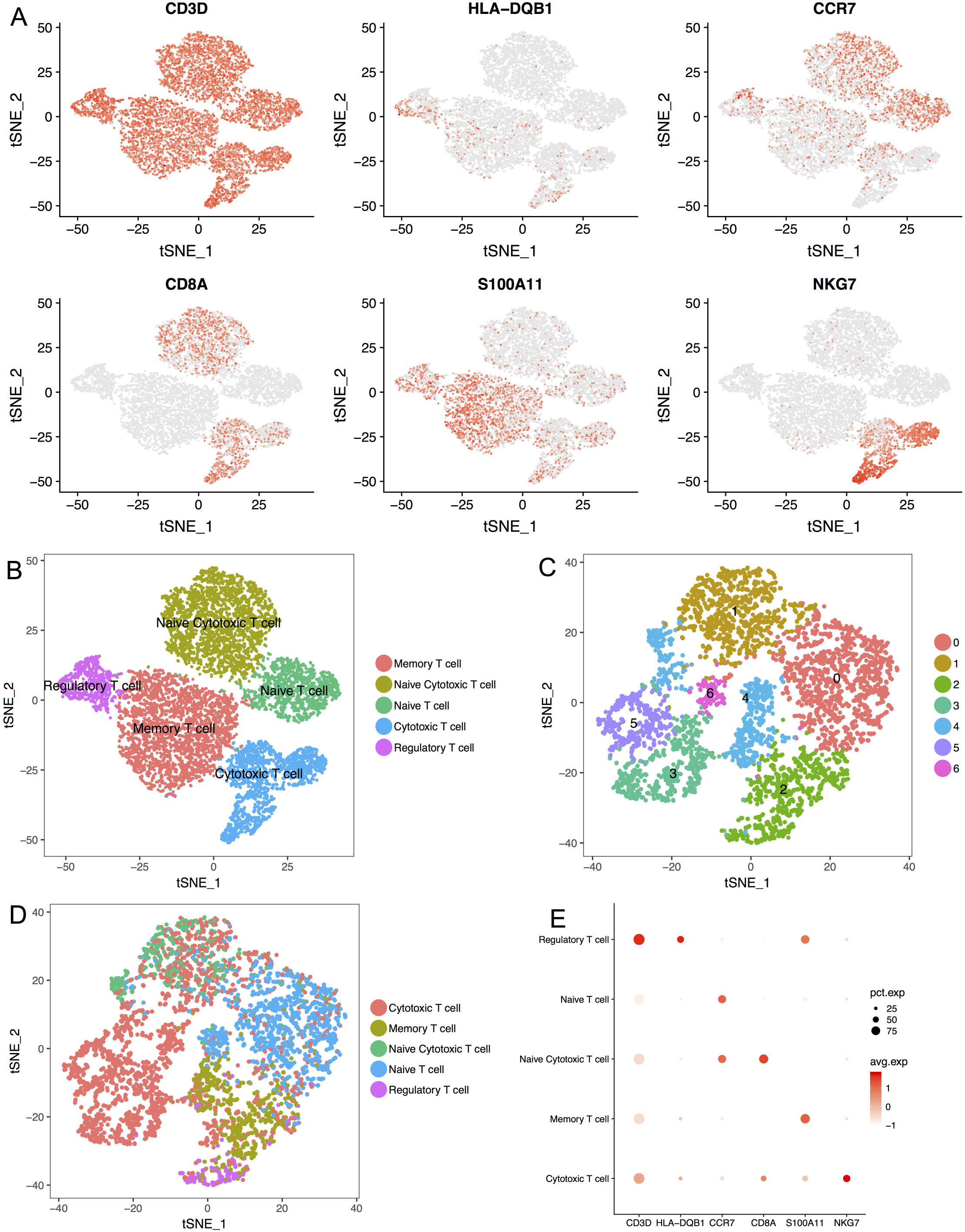
Neural network predicts sub cell types. (a) TSNE plots showing 6 maker genes’ expression for the reference T cell subtypes. (b) T cell subtypes obtained to train the neural network. (c) T cells from the human PBMC were grouped into 7 clusters by the unsupervised method. (d) Subtypes predicted for the T cells from the human PBMC. (e) Dot plot showing the expression of 6 genes for the predicted subtypes, dot size represents the percentage of cells expressing the gene, color scale represents the expression level of the gene.

To test our neural network’s ability to predict cell subtypes, we trained it on the T cell subtype reference, and predicted the subtypes for the T cells from the PBMC dataset (Fig 4D). We then identified marker genes for each predicted subtype. As expected, the marker genes matched the ones from the reference (Fig 4E). These results show that our neural network can be used to accurately identify cell subtypes. We found that the subtypes predicted by the neural network did not perfectly match the cell types associated with the Seurat clusters (Fig 4C). Some clusters contained different subtypes and some subtypes were composed of several clusters. We think the difference was influenced by two factors: 1. Unsupervised clustering considers all variance in the data, while the neural network is trained to find the difference between the subtypes; 2. It is difficult to set the parameters optimally for the unsupervised analysis, which can result in multiple cell types in one cluster or multiple clusters for one cell type.

### Discussion

scRNA-seq provides high resolution profiling of the transcriptomes of single cells. Typically, the first step in scRNA-seq analysis is to assign each cell a cell type based on our prior knowledge of marker genes. Current methods for cell type assignment first cluster the cells in an unsupervised manner and rely on the canonical markers to identify the cell types for each cluster. However, this approach has several limitations, including the fact that the clusters may not optimally segregate single cell types, and certain cell types may not have previously characterized markers. Moreover, these methods are computationally intensive, especially when the number of cells becomes large. To render cell type identification in scRNA-seq more efficient, we employed a neural network, trained it on cells with predefined cell types, and used it to predict cell types for new datasets.

We first obtained and cleaned two datasets from the Tabula Muris Consortium, then trained and tested our neural network on these datasets with or without batch effect introduced by different scRNA-seq platforms. The training accuracy always approached 100%, and the testing accuracy was around 99.8% within a platform and 99.0% when testing and training are performed across different platforms. As the cell types in the two Tabula muris atlas datasets can be mutually predicted using our neural network, we merged them and used the combined datasets as the reference to predict cell types for other datasets. The predicted cell types were well matched with the cell types assigned using the canonical markers for both the mouse and human datasets. We also trained and tested the neural network on 5 T cell subtypes and found that the predicted subtypes showed the same markers as the reference subtypes, which suggests that our neural network can be used to predict sub cell types as well.

Compared to the traditional unsupervised methods used for cell type identification, our neural network has the following advantages: 1. It uses all the genes to capture the features for each cell type instead of relying on a limited number of canonical markers. 2. It focuses the analysis on the signal associated with the variance between cell types, while unsupervised clustering tends to be affected by other sources of cell type independent variation (i.e. platform or cell cycle). 3. It requires no background knowledge of cell type markers, while the unsupervised method requires users to have prior knowledge of canonical markers for each cell type in their data. 4. It is much more computationally efficient than the traditional approach. Moreover, users can subsample the reference cells to make the computation of the neural network less compute intensive and more memory efficient.

There are some aspects of our approach that could be improved in the future. As the neural network is supervised, the quantity and quality of the reference data are critical. We anticipate that with time more cell types from larger atlases should be used to train a more comprehensive neural network. Also, better pairing of reference and test sets will undoubtedly improve performance. For example, the soon to be developed human cell atlas should be used to predict human cell types instead of the mouse cell atlas. Nonetheless, we showed that even with the current reference data our neural network is computationally efficient and accurate, and should improve cell type identification pipelines.

## Materials and Methods

### Data normalization

We used several publicly available datasets in our analyses. The mouse cell atlas datasets were collected from https://tabula-muris.ds.czbiohub.org/. The CD45 sorted leukocyte datasets were published in Winkels et al^[8]^. The T cell subtypes and PBMC datasets were collected from https://support.10xgenomics.com/single-cell-gene-expression/datasets.

To filter and normalize the data, we first identified genes that were detected in both training set and test set. The training set and the test set were then merged into one matrix based on the common genes. Next, each cell’s expression value was normalized to its total expression value and multiplied by a scale factor of 10,000. The counts were increased by 1, and the log2 value was calculated. To filter out outlier genes, the genes with the highest 1% and lowest 1% expression were removed. The gene with the highest 1% and the lowest 1% standard deviation were also removed. Finally, the matrix was split into the training set and the test set.

### Neural network configuration

We used a neural network that contains an input layer, 3 hidden layers, and an output layer. The input layer had a number of nodes equal to the number of genes in the training set. The 3 hidden layers had 100, 50 and 25 nodes, respectively. The output layer had a number of nodes equal to the number of cell types in the training set. Forward propagation was implemented as:

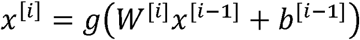

Where *x*^[*i*]^ represents the output of the *i*th layer (*x*^[0]^ represents the input layer), *b*^[*i*]^ represents the intercept of the *i*th layer, *W*^[*i*]^ represents the weight matrix of the *i*th layer, and *g* represents the activation function used in the neural network. Specifically, for the activation function, a rectified linear unit (ReLU) function was used for the input and hidden layers, which is defined as:

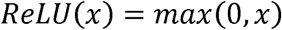

For the output layer, the softmax function was used, which is defined as:

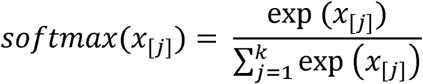

Where *x*_[*j*]_ represents the *j*th element of the input vector for the output layer, which has *k* elements, representing a total of *k* cell types in the training set.

For the loss function, we used the cross-entropy function, which is defined as:

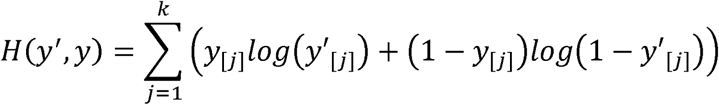

Where vector *y* represents the true label for the cell, *y*_[*j*]_ is defined to be 1 if the cell is the *j*th cell type, and the other elements in *y* are defined to be 0. *y*′ represents the output of the output layer, and *y*′_[*j*]_ represents the posterior probability that the cell is the *j*th cell type. L2 regularization was added to the loss function.

### Parameters used in the neural network

The neural network model was implemented using TensorFlow (https://www.tensorflow.org/), and the code was written in python. The parameters were initialized with Xavier initializer^[10]^. The starting learning rate was set to 0.0001 with staircase exponential decay, the decay rate was set to 0.95, and the decay step was set to 1000. This means that after every 1000 global steps, the learning rate would be the original learning rate multiplied by 0.95. 50 epochs were used to train the neural network with a mini batch size of 128, which is the number of samples used in training at every global step. The L2 regularization rate was set to 0.005.

### Unsupervised single cell analysis

To identify different cell types and find signature genes for each cell type, Seurat^[6]^ was used to analyze the digital expression matrix generated by scRNA-seq. Specifically, in Seurat, cells with less than 1000 unique molecular identifiers (UMIs) and genes detected in less than 10 cells were first filtered out. Second, highly variable genes were detected and used for further analysis. Third, the data was scaled for sequencing depth of each cell. Fourth, principle component analysis (PCA) and t-distributed stochastic neighbor embedding (tSNE) were used to reduce the dimension and plot the data on a two-dimensional graph. Lastly, a graph-based clustering approach was used to cluster the cells, then signature genes were found and used to define cell type for each cluster.

## Acknowledgments

We thank Dr. Shawn Cokus for his suggestions on early stopping of the neural network. We also thank Dr. Andrew Ng for his Deep Learning class on Coursera.

## Supporting Information

**Supplementary File 1.** This file contains 4 tables that tells the number of accurately and inaccurately predicted cells when the neural network was trained and test across the 10X and SS2 datasets.

